# The Vc2 cyclic di-GMP dependent riboswitch of *Vibrio cholerae* regulates expression of an upstream small RNA by controlling RNA stability

**DOI:** 10.1101/617340

**Authors:** Benjamin R. Pursley, Christopher M. Waters

## Abstract

Cyclic di-GMP (c-di-GMP) is a bacterial second messenger molecule that is important in the biology of *Vibrio cholerae*, but the molecular mechanisms by which this molecule regulates downstream phenotypes have not been fully characterized. We have previously shown that the Vc2 c-di-GMP-binding riboswitch, encoded upstream of the gene *tfoY*, functions as an off-switch in response to c-di-GMP. However, the mechanism by which c-di-GMP controls expression of *tfoY* has not been fully elucidated. During our studies of this mechanism, we determined that c-di-GMP binding to Vc2 also controls the abundance and stability of upstream non-coding small RNAs (sRNA) with 3’-ends located immediately downstream of the Vc2 riboswitch. Our results suggest these sRNAs are not generated by transcriptional termination but rather by preventing degradation of the upstream untranslated RNA when c-di-GMP is bound to Vc2.

**IMPORTANCE:** Riboswitches are typically RNA elements located in the 5’ untranslated region of mRNAs. They are highly structured and specifically recognize and respond to a given chemical cue to alter transcription termination or the translation initiation. In this work, we report a novel mechanism of riboswitch mediated gene regulation in *Vibrio cholerae* whereby a 3’ riboswitch, named Vc2, controls the stability of upstream untranslated RNA upon binding to its cognate ligand, the second messenger cyclic di-GMP, leading to the accumulation of previously undescribed sRNAs. We further demonstrate that binding of the ligand to the riboswitch prevents RNA degradation. As binding of riboswitches to their ligands often produces compactly structure RNA, we hypothesize this mechanism of gene regulation could be widespread.

## INTRODUCTION

*Vibrio cholerae*, the causative agent of cholera, lives in diverse environments from environmental aquatic reservoirs to microcompartments within the human host (1). The bacterial second messenger cyclic di-GMP (c-di-GMP) is a central signaling molecule in this bacterium that regulates diverse traits to enable specific adaptation to these changing conditions (2–6). *V. cholerae* encodes three know transcription factors, VpsR, VpsT, and FlrA, that bind and respond to c-di-GMP to control gene expression by regulating the transcription of genes that mediate biofilm formation, motility, Type II secretion, and DNA repair (4, 5, 7-9). In addition, *V. cholerae* also encodes two riboswitches that directly bind and respond to changes in the intracellular concentration of c-di-GMP, although the function of these riboswitches in the adaptation of *V. cholerae* is not well understood (10, 11).

C-di-GMP-binding riboswitches can be found as two distinct structural classes (12). The riboswitches encoded by *Vibrio cholerae*, named Vc1 and Vc2, are both designated as class I c-di-GMP-binding riboswitches (10, 11). Vc1 is encoded in the 5’-untranslated region (UTR) of the putative adhesin *gbpA*, and binding of c-di-GMP to Vc1 enhances production of GbpA, making this riboswitch an “on-switch” (11). Vc2, which has extensive sequence identity with Vc1, is encoded in the 5’-UTR of the *tfoY* gene (13). We recently demonstrated that Vc2 functions as an off-switch as it represses TfoY production when bound to c-di-GMP, leading to a decrease in TfoY-mediated dispersive motility (10). Other bacterial species also encode cyclic di-nucleotide binding riboswitches. For example, *Clostridium difficile* encodes 11 class I and 4 class II c-di-GMP-binding riboswitches (14). A recent analysis in *C. difficile* suggested that the class I riboswitches are primarily off-switches while the class II riboswitches are on-switches (14). *Geobacter sulfurreducens* encodes riboswitches that can discriminate between c-di-GMP and bacterial cyclic GMP-AMP, and these riboswitches appear to regulate genes involved in exoelectrogenesis (15, 16).

Our previous studies showed that the regulation of the *tfoY* gene is complex consisting of four promoters we have designated *P_1-4-tfoY_* and the Vc2 c-di-GMP binding riboswitch (Fig. 1A)(10). This regulatory system discriminates between at least three distinct intracellular c-di-GMP concentrations in *V. cholerae* when grown in rich medium in the laboratory: low, generated by overexpressing an active EAL; intermediate, the unaltered wild type levels of c-di-GMP; or high, generated by overexpressing an active DGC. Importantly, we demonstrated that the concentrations of c-di-GMP observed in these three states is physiologically relevant (10). Transcription from the promoters *P_3-tfoY_* and *P_4-tfoY_*, which are encoded within and to the 3’ end of Vc2, respectively, and located closest to the translation start site of *tfoY*, do not lead to transcription of a complete Vc2 riboswitch, and thus expression of *tfoY* from these transcripts is Vc2-independent (Fig. 1A. Rather, these promoters are induced in a VpsR-dependent manner at high intracellular c-di-GMP. Transcription from the upstream promoters, *P_1-tfoY_* and *P_2-tfoY_*, does generate the full length Vc2 riboswitch, and we demonstrated binding of c-di-GMP to Vc2 leads to inhibition of TfoY production at intermediate and high c-di-GMP concentrations via an unknown mechanism (10).

**Figure 1:**
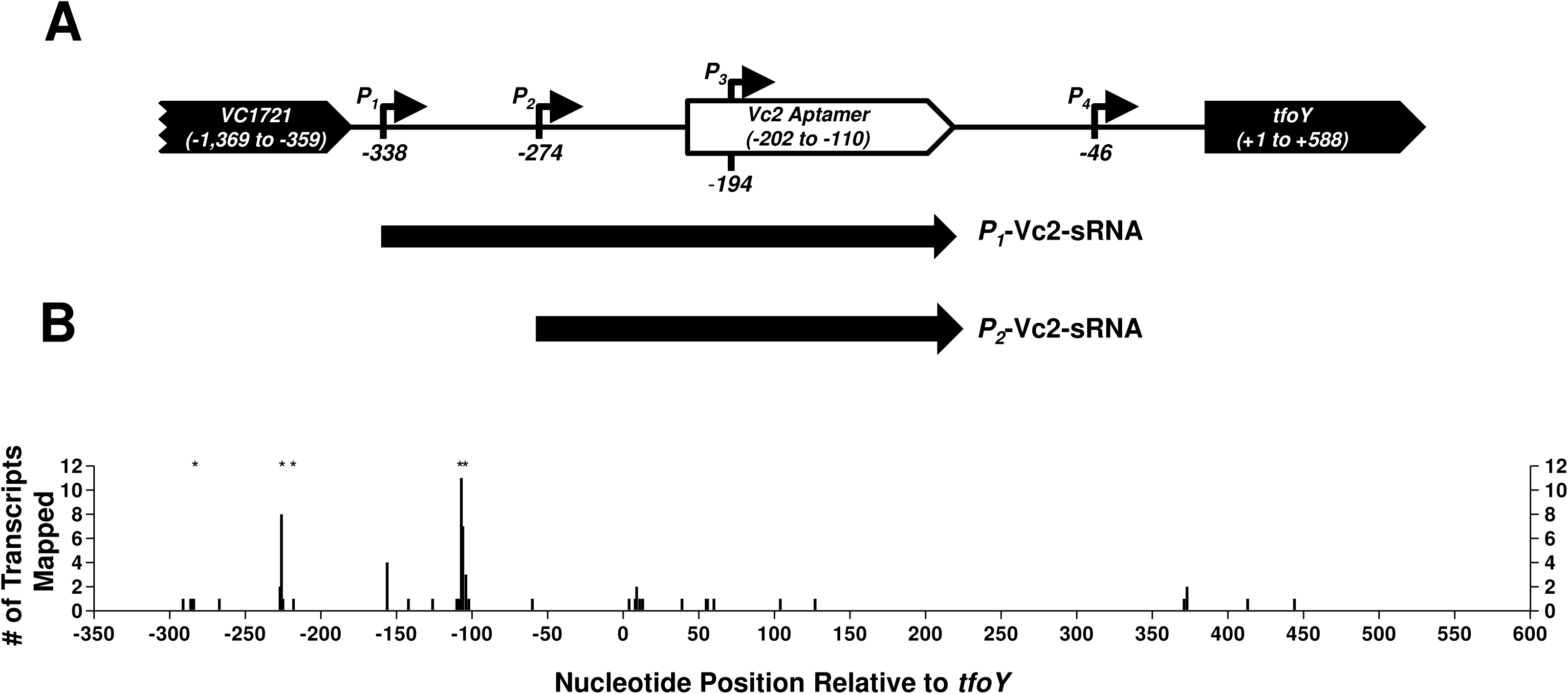
3’-R.A.C.E. of *P_1-tfoY_* transcripts. **A)** This map shows the four transcriptional start sites identified for *tfoY* transcripts, and their location on *V. cholerae* chromosome. Numbering is relative to the start of the *tfoY* coding sequence. The location of the sRNAs identified in this study are indicated below the map. **B)** Numbering of the X-axis corresponds to the genomic map in 1A, except for position zero, which is included here for reference but does not represent a nucleotide position on the genome. The Y-axis indicates the number of *P_1-tfoY_* transcript sequences recovered whose 3’-end mapped to the position indicated. Asterisks indicate sites where sequences contained 3’-end tails with extra-genomic nucleotides. 66 sequences are depicted in total.

These studies suggested a model whereby one function of Vc2 was to inhibit production of TfoY from the upstream *P_1-tfoY_* and *P_2-tfoY_* at intermediate and high concentration of c-di-GMP while transcription initiation of *tfoY* from *P_3-tfoY_* and *P_4-tfoY_* at high c-di-GMP concentrations was induced by c-di-GMP binding to the transcription factor VpsR. At low concentrations of c-di-GMP, TfoY stimulates a rapid, flagellar-based swimming motility that we termed dispersive motility (10), but the function of increased TfoY at high c-di-GMP concentrations is not known. Both *in vitro* and *in vivo* evidence suggested that Vc2 inhibits translation of *tfoY* when bound to c-di-GMP with the ribosome binding site (RBS) of *tfoY* sequestered by an anti-RBS element located downstream of the Vc2 riboswitch (17, 18), but the mechanism of this inhibition has not yet been fully established in the native sequence of the *tfoY* 5’-UTR.

While further exploring the mechanism by which Vc2 functions as an off-switch, we observed that most of the 3’-ends of transcripts derived from the *P_1-tfoY_* promoter were located at the 3’-end of the Vc2 riboswitch, leading us to elucidate that c-di-GMP binding to the Vc2 riboswitch directs accumulation of a novel sRNA. This sRNA, which we named the *P_1_*-Vc2 sRNA, initiates at the upstream *P_1-tfoY_* promoter and has a 3’-end located at end of the Vc2 riboswitch. The *P_1_*-Vc2 sRNA is not generated by transcription termination, but rather it accumulates at intermediate and high intracellular c-di-GMP concentrations due to increased stability when Vc2 is bound to c-di-GMP. Our results describe a novel form of gene regulation whereby a 3’-riboswitch controls the accumulation of an untranslated sRNA by preventing its degradation.

## RESULTS

### Identification of sRNAs that contain Vc2 on their 3’-end

In our previous study, primer-extension analyses determined that the majority of transcripts containing the *tfoY* open reading frame (ORF) originate from *P_3-tfoY_* and *P_4-tfoY_*, even though *P_1-tfoY_* and *P_2-tfoY_* appeared to be stronger promoters based on expression analysis of *gfp* transcriptional fusions (10). Therefore, the fate of transcripts initiated at *P_1-tfoY_* and *P_2-tfoY_* was unclear. To further explore this finding, we conducted a 3’-Rapid Amplification of cDNA Ends (RACE) assay to identify the location of the 3’-ends of transcripts originating from *P_1-tfoY_*. These studies were performed with wild type *V. cholerae* and thus were at intermediate concentrations of c-di-GMP. At these concentrations, c-di-GMP is bound to Vc2 and TfoY production is reduced (10). Prior studies had predicted that Vc2 regulates translation of *tfoY* (17, 18), leading us to hypothesize that the 3’ ends of *P_1-tfoY_* transcripts would be located downstream of the *tfoY* gene. In total, we recovered 66 sequences which mapped to the *VC1721-tfoY* intergenic region (Fig. 1B). We did not recover any transcripts with a 3’-end mapping downstream of the *tfoY* stop codon; however, we did recover multiple transcripts whose 3’-ends were dispersed within the *tfoY* coding sequence, suggesting high amounts of mRNA degradation.

Surprisingly, the 3’-end of the majority of sequences was located upstream of the *tfoY* translational start site. The greatest number of 3’-ends we observed, 38% of all recovered transcripts, mapped to the region −110 to −102 relative to the start of the *tfoY* coding sequence, which is the region immediately adjacent to the 3’-end of the P1 stem-loop that forms the base of the Vc2 riboswitch (19). An additional 17% of the sequences contained a 3’-end located further upstream, between −227 to −225 relative to the start of the *tfoY* coding sequence. This region correlates with the 3’-end of a putative *rho*-independent terminator located between the *P_2-tfoY_* and *P_3-tfoY_* transcriptional start sites, and it is also the location of the identified VpsR binding sites (10, 20). Of all the transcripts recovered, 30% featured 3’-end tails containing between 1 and 4 extragenomic nucleotides, almost exclusively adenines, and the majority of these tailed transcripts mapped to the aforementioned sites of the Vc2 aptamer or the upstream *rho*-independent terminator. These results led us to hypothesize that most of the transcripts initiated from *P_1-tfoY_* and *P_2-tfoY_* exist as sRNAs with the Vc2 riboswitch aptamer located at their 3’ ends, and these transcripts did not contain a full-length *tfoY* sequence, explaining the results of our previous primer extension assay.

### Multiple sRNAs are transcribed from the *VC1721-tfoY* intergenic region

To confirm that the sRNA species predicted by our 3’-RACE analysis were produced by the upstream *tfoY* promoters in *V. cholerae*, we performed a series of Northern blots with probes encompassing different segments of the *VC1721-tfoY* intergenic region against RNA harvested from WT *V. cholerae* (Fig. 2). Four probes were used that are complementary to the following regions: probe *P_1_*(−337 to −272, between the *P_1-tfoY_* and *P_2-tfoY_* start sites), probe *P_2_* (−272 to −191, between the *P_2-tfoY_* and *P_3-tfoY_* start sites), probe *P_1+2_* (−337 to −191, a larger sequence overlapping the *P_1-tfoY_* to *P_3-tfoY_* start sites), and probe Vc2 (−211 to −94, the sequence of the Vc2 riboswitch).

**Figure 2:**
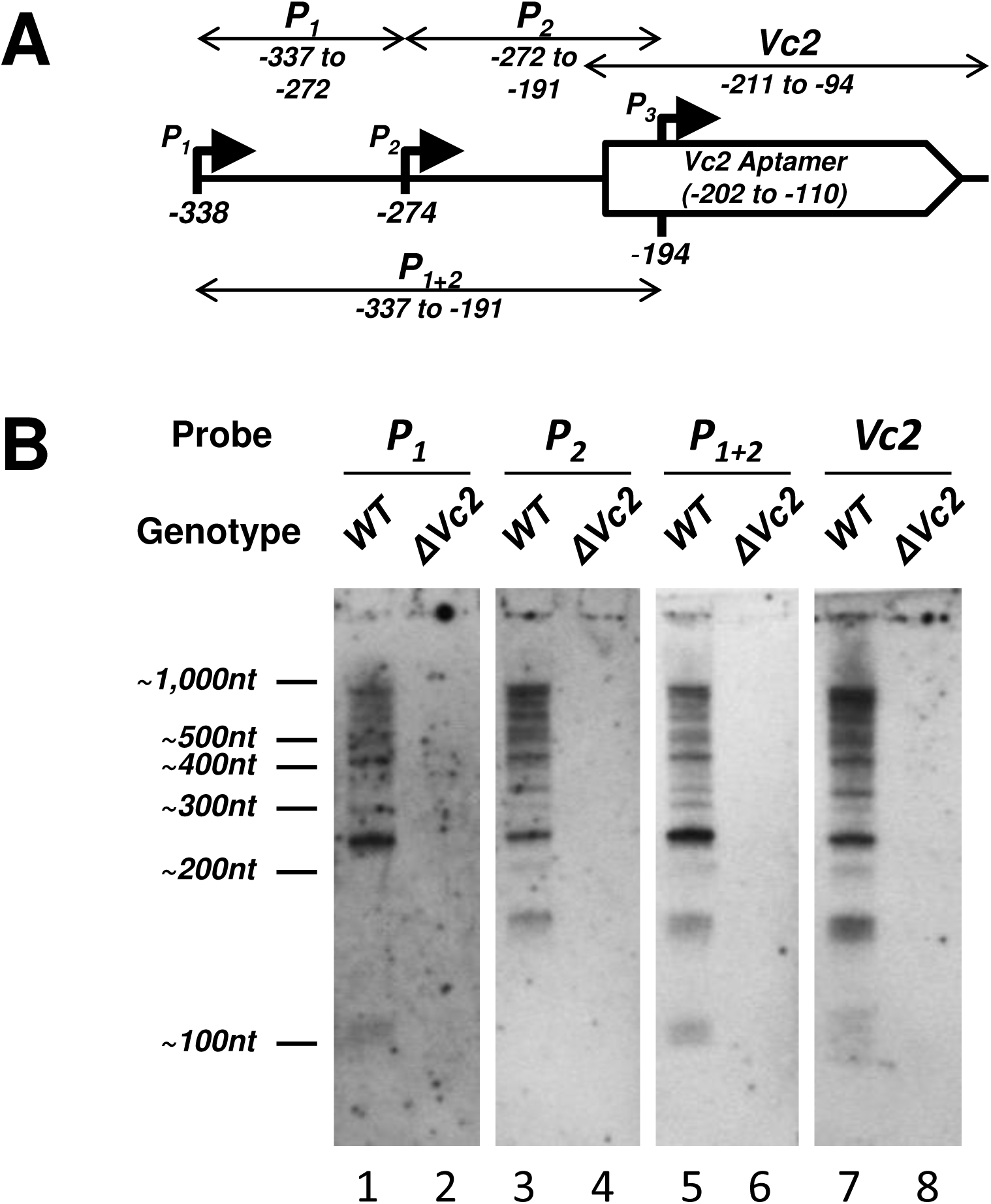
Northern Blot Analysis of *VC1721-tfoY* Intergenic Region. **A)** The location of the four RNA probes used for the Northern blot in Fig. 2B and Fig 3 is shown. **B)** Northern blot analysis of RNA extracted from either WT or the DVc2 mutant *V. cholerae*. The location of the ladder bands is shown to the left of the blots.

Based on the results of the 3’-RACE assay, we expected to observe specific sRNA transcripts of ∼231 nucleotides and ∼167 nucleotides in size originating from the *P_1-tfoY_* and *P_2-tfoY_* promoters, respectively, with both ending at the immediate 3’ edge of the Vc2 aptamer. The presence of the ∼231 nucleotide band in all four blots (Fig. 2B, Lanes 1, 3, 5, and 7), and the presence of the ∼167 nucleotide band in all but the probe *P_1_* blot (Fig. 2, Lanes 3, 5, and 7), confirms the location of these sRNAs species relative to the promoters and the riboswitch aptamer. We refer to these ∼231 nucleotide and ∼167 nucleotide transcripts henceforth as the *P_1_-Vc2* and *P_2_-Vc2* sRNAs, respectively. We also expected to observe a sRNA ∼112 nucleotides in size initiated from the *P_1-tfoY_* promoter and ending at the predicted intrinsic terminator immediately downstream of the *P_2-tfoY_* start site. The appearance of this band in the probe *P_1_* and *P_1+2_* blots and its absence from the probe *P_2_* blot confirmed the presence of this transcript as well.

All of the probes detected a large number of longer RNA species extending into the *tfoY* coding sequence. This is consistent with our 3’-RACE results identifying multiple 3’ RNA ends in this region (Fig. 1), and it further suggests that transcription from P*_1-tfoY_* and P*_2-tfoY_* can extend into the *tfoY* ORF. All the signals in the Northern blot analysis from every probe used were lost in the ΔVc2 mutant, which has a deletion from −359 to −43 on the genome of *V. cholerae*, showing that these RNA species are specific to the Vc2 riboswitch locus. It is also worth noting that the coding sequence of *VC1721*, the nearest gene upstream of the riboswitch, is 1,011 nucleotides in length. Therefore, based on their size, all the transcripts detected here should have originated downstream from the *VC1721* ORF and do not represent run-off transcription from upstream *VC1721* mRNA. These data show that the *tfoY* upstream region produces multiple sRNA species. We name these the *P_1_*-Vc2-sRNA and *P_2_*-Vc2-sRNA for the promoter from which they are transcribed.

### The Vc2 riboswitch and c-di-GMP regulate the abundance of sRNAs

To investigate the effect of changes in c-di-GMP on Vc2 riboswitch-containing transcripts *in vivo*, we performed a higher resolution Northern blot of *V. cholerae* RNA extracts with probe Vc2 (Fig. 3). An initial observation across all of our samples is that there are many different Vc2 riboswitch-containing transcripts between 300 and 1000 nucleotides in length (Fig. 3). The annotated coding sequence of *tfoY* is 588 nucleotides, therefore the minimum size of full-length *tfoY* mRNAs should be 926 nucleotides if originating from *P_1-tfoY_*, or 782 nucleotides if originating from *P_3-tfoY_*. This implies that a significant number of the transcripts detected contain only a fraction of the annotated *tfoY* ORF, again consistent with data from our 3’-RACE analysis (Fig. 1). An integrity analysis of ribosomal RNA from our samples using an Agilent Bioanalyzer confirmed that our RNA was of high quality with little to no degradation occurring during extraction. From this we conclude that most Vc2 riboswitch-containing transcripts are inherently unstable and subject to a high rate of degradation *in vivo*.

**Figure 3:**
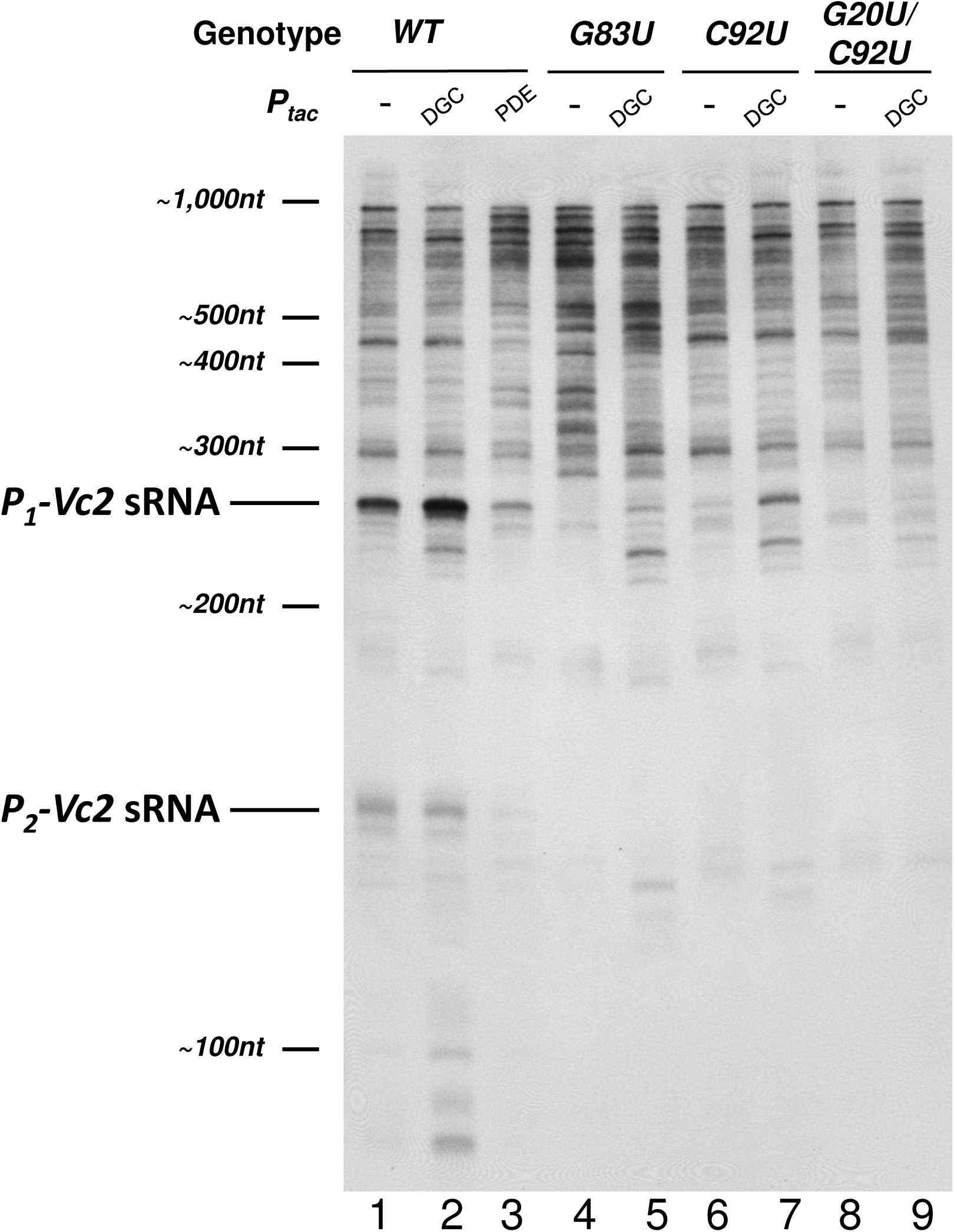
Northern Blot Analysis of *V. cholerae* Riboswitch Mutants. Lanes are as described in the main text. This figure is a Northern blot of the four *V. cholerae* strains indicated at the top of the blot with the Vc2 probe showin in Fig. 2A. It is representative of multiple experiments. The film shown here is overexposed in order to aid the visualization of bands. Values reported in the text were quantified from blots exposed to film over a linear range of detection. DGC=overexpression of QrgB, PDE=overexpression of VC1086.

Regardless of the transcriptional processes occurring downstream of Vc2, under wild-type conditions the *P_1_-Vc2* sRNA is the most abundant single RNA species produced that contains the Vc2 sequence (Fig. 3, Lane 1). This is consistent with the disparity we had previously observed between the high activity of the *P_1-tfoY_* promoter and its low contribution to total *tfoY* mRNA (10). To determine if this sRNA is impacted by changing c-di-GMP levels, we expressed from plasmids the diguanylate cyclase (DGC) enzyme QrgB to increase c-di-GMP concentrations or the phosphodiesterase (PDE) enzyme VC1086 to decrease c-di-GMP concentrations. We have previously demonstrated these enzymes generate physiologically relevant concentrations of c-di-GMP in *V. cholerae* (10). Upon DGC induction, we observed a 2.5-fold increase in the abundance of the *P_1_*-Vc2 sRNA (Fig. 3, lane 2). Our previous results show that the transcription of *P_1-tfoY_* is not significantly affected by induction of QrgB (10), implying that the increase in *P_1_*-Vc2 sRNA observed here is the result of a post-transcriptional mechanism. On the other hand, induction of the PDE, which decreased c-di-GMP, led to decreased abundance of the *P_1_*-Vc2 sRNA (Fig. 3, lane 3).

The *P_2_*-Vc2 sRNA that we previously observed was also evident as a 167-nucleotide band, although it was not as abundant as the *P_1_*-Vc2 sRNA. The abundance of the *P_2_*-Vc2 sRNA was slightly decreased upon DGC induction (Fig. 3, lane 2). This decrease was expected and can be attributed to c-di-GMP-mediated repression of the *P_2-tfoY_* promoter that we have previously reported (10). The *P_2_*-Vc2 sRNA was nearly lost upon induction of the PDE (Fig. 3, lane 4)

We hypothesized that the post-transcriptional mechanism responsible for the increase in *P_1_*-Vc2 sRNAs during changes in c-di-GMP concentrations is dependent on c-di-GMP binding to the Vc2 aptamer. To test this, we generated a number of chromosomal *V. cholerae* mutations in Vc2 to assess its function. We first constructed a G83U chromosomal mutation in the Vc2 riboswitch as this has been shown to disrupt its structure. When this mutation is present, the *P_1_*-Vc2 sRNA is not visible at wild type levels of c-di-GMP and only very faint when the DGC is induced (Fig. 3, Lanes 4 and 5).

We next mutated the C92 and G20 bases in Vc2 to uracils on the *V. cholerae* chromosome to assess the role of c-di-GMP binding in formation of these sRNAs. The G20 and C92 bases of the Vc2 riboswitch are the sites which directly interact with the guanosine bases of c-di-GMP, and substitutions of uracil at either of these sites significantly disrupts ligand binding without general altering the ligand-free structure (19, 21). Accordingly, in strains with either single or double mutations at the ligand binding sites we observed loss of the *P_1_*-Vc2 sRNA (Fig. 3, Lanes 6 and 8). In the C92U mutant, the reduction in abundance from wild-type was 30-fold, and in the G20U/C92U mutant the amount of *P_1_*-Vc2 sRNA remaining was below the limit of detection. We also observed complete loss of the *P_2_*-Vc2 sRNA in both of these mutant backgrounds. Expression of the DGC QrgB in the C92U and G20U/C92U strains led to partial recovery of the abundance of the *P_1_*-Vc2 sRNA in the C92U mutant but virtually no recovery in the G20U/C92U double mutant (Fig. 3, Lanes 7 and 9). The relative strength of this recovery in each strain was consistent with the previous biochemical data that the single binding site mutant retains a higher c-di-GMP affinity than the double mutant (19).

### The Vc2 riboswitch is not sufficient for transcription termination *in vitro*

The observation that short transcripts are produced which contain the Vc2 aptamer at the 3’-end could be explained if the expression platform of the Vc2 riboswitch functions as a *rho*-independent transcriptional terminator. No readily apparent terminator structure is located adjacent to the Vc2 riboswitch, and bioinformatic analysis with *rho*-independent terminator prediction software indicates that the best putative terminator nearby is located at the +193 to +235 region of the *tfoY* coding sequence, which is more than 300 nucleotides downstream of the 3’-end of the Vc2 aptamer (13, 20, 22). Furthermore, recent publications suggested that Vc2 regulates *tfoY* by controlling translation, not transcription (17, 18). Nevertheless, to experimentally test if the Vc2 aptamer stimulates transcription termination upon binding to c-di-GMP, we conducted an *in vitro* transcription termination assay with *E. coli* RNA polymerase complexed with σ^70^ using a linear PCR template encompassing the genomic region −535 to +120 relative to the *tfoY* coding sequence.

Because our transcription template included all of the *tfoY* promoters, we expected to see full-length transcription products of sizes 458, 394, 314, and 166 nucleotides corresponding to the observed transcriptional start sites of the *P_1-tfoY_*, *P_2-tfoY_*, *P_3-tfoY_*, and *P_4-tfoY_* promoters, respectively, and indeed all of those transcripts were observed (Fig. 4, Lane 1). Only two other major transcripts were detected, one >600 nucleotides, likely generated from end-to-end transcription of the template, and one at the ∼180 nucleotides size range. Most notably, we did not observe any major transcript in the size range of the *P_1_*-Vc2 sRNA, which is the most prominent riboswitch-containing transcript generated *in vivo*.

**Figure 4:**
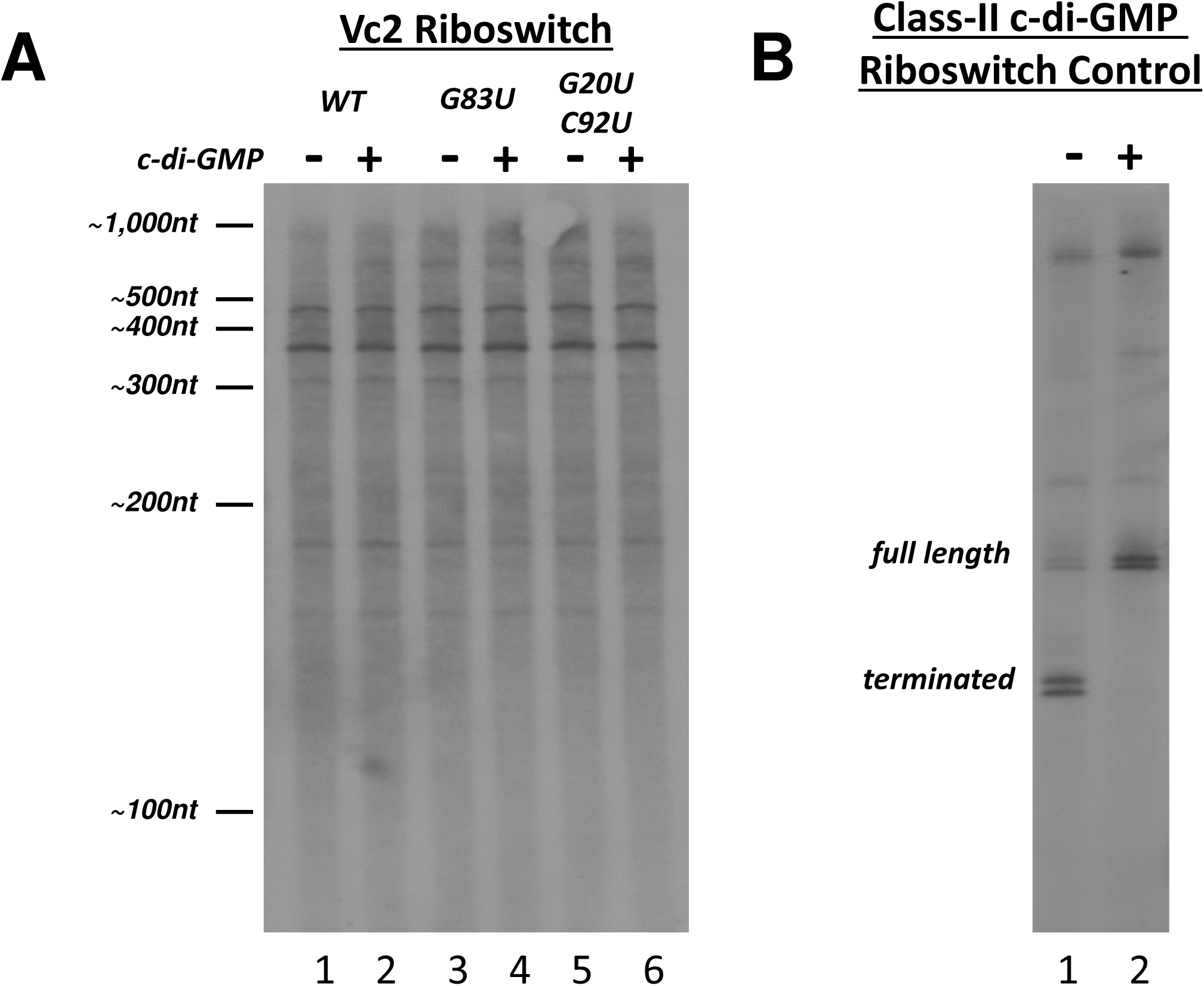
*in vitro* Termination Assay with c-di-GMP. **A)** An *in vitro* transcription reaction with *E. coli* RNA polymerase and sigma 70 was performed on a DNA template containing the sequence upstream of *tfoY* and part of the *tfoY* open reading frame. Mutations to the riboswitch is indicated. **B)** A similar *in vitro* transcription reaction was performed with a class II riboswitch from *Clostridium difficile*. C-di-GMP was added at a concentration of 0.1 mM.

We tested the possibility that Vc2 riboswitch could induce termination by adding exogenous c-di-GMP to the transcription reaction (Fig. 4, Lane 2), yet no changes in the size or abundance of transcripts were observed. We also tested the effect of disrupting the structural integrity of the riboswitch by using transcription templates containing the same mutations in the aptamer sequence which had earlier been investigated *in vivo*. Neither a G83U structural mutation that disrupts tertiary structure of the riboswitch nor the G20U/C92U binding site mutations described earlier had any detectable effect on the production of transcripts *in vitro*. Therefore, the riboswitch had no impact on transcription of this region *in vitro* with or without c-di-GMP.

To ensure that our reaction conditions were sufficient for detecting potential riboswitch mediated termination, we tested a transcriptional template comprising the *ompR*-associated class-II c-di-GMP riboswitch of *Clostridium difficile*, for which a transcription termination mechanism had previously been demonstrated *in vitro* (23). For this control experiment, addition of c-di-GMP inhibited transcription termination, consistent with the known mechanism of regulation of this riboswitch (Fig. 4B). Since we observed c-di-GMP regulation of termination with the control riboswitch but not with the Vc2 riboswitch, we conclude that the Vc2 riboswitch does not generate the *P_1_*-Vc2 and *P_2_*-Vc2 sRNAs through *rho*-independent termination.

### Stability of the P_1_-Vc2 sRNA is c-di-GMP dependent

Because Vc2 does not appear to regulate transcription termination and we observe significant RNA degradation in this region, we hypothesized that the Vc2 aptamer controls the post-transcriptional stability of the Vc2 sRNAs. To test this hypothesis, we added rifampicin to halt transcription at mid-log phase growth of *V. cholerae* cells under conditions of both wild-type and elevated c-di-GMP, and subsequently extracted RNA from the cells over a series of multiple time intervals (Fig. 5). These experiments were not informative when a PDE was expressed or with the Vc2 mutants as these conditions had too low of levels of the *P_1_-* Vc2 sRNA to perform this analysis. Transcripts containing the Vc2 aptamer sequence were detected by Northern blot with probe Vc2, and we then quantified the amount of *P_1_-* Vc2 sRNA remaining at each time point after the addition of rifampicin relative to the amount of *P_1_-* Vc2 sRNA at time zero (Fig. 5B).

**Figure 5:**
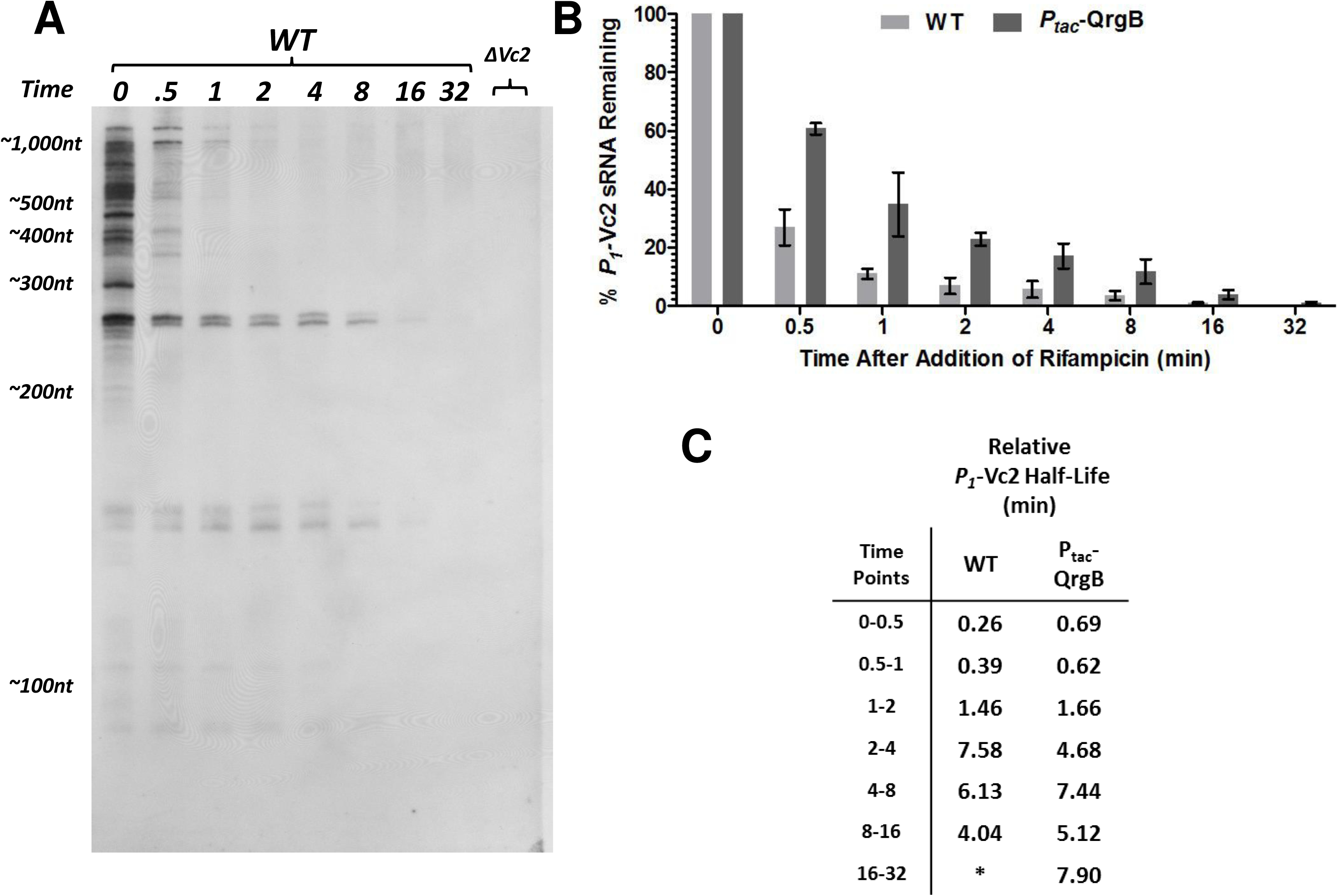
RNA Stability Assay with Rifampicin. **A)** A single exposure of a Northern blot from one of the replicates of the RNA stability assay. Time is indicated as minutes after rifampicin treatment. Measurements of RNA abundance were performed from a series of exposures in order to quantify the intensity of each band on the blot within its appropriate liner range. **B)** Stability assays were performed in triplicate for cells grown under both wild-type and elevated c-di-GMP conditions and the mean percent of *P_1_*-Vc2 sRNA transcript remaining at each time point is depicted. Error bars indicate the standard deviation among three biological replicates. **C)** From the same data shown in A, the half-life of the *P_1_*-Vc2 sRNA was calculated for the period between each time point.

Our analysis revealed that the *P_1_*-Vc2 and *P_2_*-Vc2 sRNA species are significantly more stable than all other transcripts originating from the *VC1721-tfoY* intergenic locus (Fig. 5A). Surprisingly, this analysis also revealed that the *P_1_*-Vc2 and *P_2_*-Vc2 sRNAs are not single uniform RNA species, but instead are both represented by a doublet of transcripts with a 5 to 10 nucleotide difference in size (Fig. 5A). Although it was not determined in this specific assay, these transcripts may represent the most prominent 3’-tailed and non-tailed sRNA species we observed in our 3’-RACE experiments. For the purposes of our quantitative analysis of the amount of transcript remaining, both bands at the *P_1_*-Vc2 sRNA size range were quantified together as the total amount of *P_1_*-Vc2 sRNAs.

Our RNA stability assay indicated that the individual members of the *P_1_*-Vc2 sRNA population, on average, persist for longer in the cell when c-di-GMP is elevated (Fig. 5B). This difference in stability between conditions of low and high c-di-GMP is a sufficient mechanism to explain the difference in the abundance of the *P_1_*-Vc2 sRNA between those same conditions. Interestingly, the observed degradation of the *P_1_*-Vc2 sRNA does not adhere to a simple one-phase decay model. Instead, this sRNA appears to be degraded rapidly at early time points and slowly at later time points. To compare the stability of the *P_1_*-Vc2 sRNA transcripts over time, we calculated the effective half-lives observed for the transcripts relative to each set of time points (Fig. 5C). At the first two time points in wild-type conditions, the half-life of the *P_1_*-Vc2 sRNA is less than half a minute, while at later time points this half-life has risen to the range of 4 to 8 minutes. Under elevated c-di-GMP conditions, the half-life of the *P_1_*-Vc2 sRNA starts at more than twice the length of the wild-type conditions, but then settles into a similar range of 4 to 8 minutes (Fig. 5C).

The observed pattern of increasing half-lives is consistent with there being multiple versions of the *P_1_*-Vc2 sRNA with different rates of decay. Because the *P_1_*-Vc2 sRNA contains a functional Vc2 aptamer domain, we hypothesize these different versions of transcript to be c-di-GMP-bound and -unbound forms of the *P_1_*-Vc2 sRNA. In this way, the average half-life of the *P_1_*-Vc2 sRNA is changing over time because at early time points many members of the *P_1_*-Vc2 sRNA population do not contain a ligand-bound riboswitch and are rapidly degraded, but at later time points all the unbound transcripts have been degraded and only stable ligand-bound *P_1_*-Vc2 sRNAs remain. From this interpretation, the 4 to 8-minute half-life measurement at later time points represents the closest approximation to the actual half-life of the c-di-GMP-bound form of the *P_1_*-Vc2 sRNA, and the actual half-life of the c-di-GMP-unbound form of the *P_1_*-Vc2 sRNA is less than the lowest measured half-life at the earliest time point, i.e. less than 15.6 seconds. When compared to studies on global RNA stability in other bacteria, this shift in the stability of the *P_1_*-Vc2 sRNA is quite dramatic, as measurements in *E. coli* indicate the half-life of most RNAs to be in the range of 2 to 8 minutes (24–27).

## DISCUSSION

In this report, we describe a novel regulatory mechanism whereby biding of a ligand to a 3’-riboswitch regulates the stability of the upstream sRNA. Our findings arose from the observation that the vast majority of Vc2 riboswitch-containing transcripts do not ultimately contain *tfoY* mRNA. Instead, the upstream *tfoY* promoters produce two major sRNA species, the most prominent of which, designated as the *P_1_*-Vc2 sRNA, originates at the *P_1-tfoY_* promoter and features the Vc2 riboswitch at its 3’ terminus. Our results are consistent with the results of multiple transcriptomic studies in *V. cholerae* which have reported numerous sRNAs that map to the *VC1721-tfoY* intergenic region (Fig. S1) (28–32).

Superficially, the observation that increased c-di-GMP leads to production of short transcripts gives an impression that the Vc2 riboswitch serves as a transcriptional terminator. However, our results do not support that conclusion as we did not observe transcription termination mediated by the Vc2 riboswitch *in vitro*. Furthermore, termination at a site downstream of the riboswitch would not explain how the *P_1_*-Vc2 sRNA becomes truncated at the immediate 3’-end of the P1 stem that forms the base of the Vc2 aptamer structure. Riboswitches that function by intrinsic termination employ an additional RNA structure, termed the expression platform, which is adjacent to, but separate from, the aptamer domain (33). For the Vc2 riboswitch, no such structure has been identified downstream of the aptamer domain (13, 22). Finally, both primer extension analysis and Northern blotting indicates that transcription initiated at the *P_1-tfoY_* promoter is regularly able to proceed past the riboswitch (10). No evidence, therefore, supports the hypothesis that the Vc2 riboswitch functions by controlling transcription termination.

We previously demonstrated that Vc2 bound to c-di-GMP inhibits production of TfoY leading to decreased motility (10). Here, we demonstrate that a second phenotype for Vc2 is the changing abundance of the *P_1_*-Vc2 and *P_2_*-Vc2 sRNAs. When intracellular c-di-GMP concentrations are elevated, more *P_1_*-Vc2 sRNA is present, and mutation of Vc2 to disrupt its structure or ability to bind c-di-GMP leads to less *P_1_*-Vc2 sRNA. The *P_2_*-Vc2 sRNA does not increase at higher concentrations of c-di-GMP due to repression of transcription of the *P_2-tfoY_* promoter (10), but its stability is still dependent on Vc2 binding to c-di-GMP. Because the activity of the *P_1-tfoY_* promoter is not significantly affected by changes in c-di-GMP, the changing abundance of the *P_1_*-Vc2 sRNA must be the consequence of post-transcriptional regulation. Our data indicate that the c-di-GMP-unbound *P_1_*-Vc2 sRNA is subject to very rapid turnover by the RNA degradosome, but the c-di-GMP-bound *P_1_*-Vc2 sRNA is significantly more resistant to degradation. Thus, we conclude that the Vc2 riboswitch functions to control the cellular levels of the upstream RNA by ligand-mediated aptamer protection from 3’ degradation. Considering that a characteristic of aptamer domains is to fold into a compact nucleotide structure around the target ligand we expect that this form of riboswitch mediated gene regulation may be widespread.

Riboswitches typically consist of two elements, a ligand binding aptamer domain and a cis-encoded expression platform that differentially interacts with the aptamer based on whether it is bound to its cognate ligand. Two important papers have concluded that Vc2 regulates *tfoY* by altering translation (17, 34). These authors proposed a structural model in which the P1 stem loop that is formed upon binding of c-di-GMP to Vc2 leads to sequestration of the *tfoY* RBS. Alternatively, when c-di-GMP concentrations are low, the P1 loop is not formed but rather interacts with downstream sequence to prevent formation of the anti-RBS structure. Although these structural models are not yet fully validated in the native Vc2 sequence, it seems clear that binding of c-di-GMP to Vc2 impacts formation of a downstream expression platform to inhibit translation. However, when considering the upstream sequence, our data suggest that the Vc2 riboswitch prevents its degradation by stabilizing the 3’-ends of sRNAs. This raises the intriguing possibility that for the upstream *P_1_*-Vc2 sRNA and *P_2_*-Vc2 sRNA the aptamer domain itself might dually function as the expression platform. It is also possible that sequences located 3’ of Vc2 that are impacted by c-di-GMP binding could be important in regulating RNA degradation.

Our model of how the Vc2 riboswitch regulates RNA processing is unique from known mechanisms of riboswitch-mediated RNA degradation. The best characterized examples of riboswitches that both produce small RNA fragments and regulate RNA processing include the *glmS* riboswitch and ribozyme, the *yitJ* riboswitch and RNase Y, and the *lysC* riboswitch and RNase E (35–37). In all those systems the effect of ligand-binding is destabilizing because it promotes degradation of riboswitch-containing transcripts. The Vc2 riboswitch differs from those previous examples as c-di-GMP binding inhibits degradation of the upstream sRNAs. The only previous example of 3’-end riboswitch-regulated RNA degradation comes from the plant domain. In that case, a thiamin pyrophosphate riboswitch that is located downstream of a thiamin biosynthetic gene controls RNA processing of the 3’-UTR in a manner that determines the expression level of the mature mRNA (38). But again, whether ligand bound or not, the structure of the riboswitch aptamer is sacrificed in either of the processing outcomes and the mature transcript does not contain a functional riboswitch (38).

Likewise, only a handful of riboswitch-containing sRNAs have been described. An *S*-adenosylmethionine riboswitch in *Listeria monocytogenes* induces transcription termination upon ligand binding, and then upon discarding the ligand the bases of the aptamer rearrange and interact through complementarity with a target mRNA (39). Two B12-binding riboswitches regulate non-coding RNAs in *Listeria monocytogenes*. A riboswitch located 3’ and antisense to the *pocR* gene controls the termination of the counter-transcribed RNA AspocR such that in the absence of B12 full length AspocR is produced inhibiting expression of *pocR* (40). In addition, a B_12_-binding riboswitch regulates the length of the non-coding *ril55* sRNA in *L. monocytogenes* by controlling a factor dependent-transcriptional terminator, which interestingly regulates the downstream gene by binding to a response regulator (41). A type-I c-di-GMP-binding riboswitch in *Bdellovibrio bacteriovorus* was found to be more highly represented in an RNA-Seq data set than all other non-rRNA, non-tRNA transcripts, and was hypothesized to function as a storage bank for c-di-GMP that controls the transition between the growth and attack phases of the cell although this hypothesis was never directly tested (42). A highly abundant sRNA in *Rhodobacter sphaeroides* is derived from a 5’-UTR region that encoded a potential riboswitch, although the function of the riboswitch nor its role in regulating the sRNA was never shown (43). Likewise, an RNA-Seq study focusing on regulatory RNAs in *C. difficile* included an analysis of the expression of genes downstream of various riboswitches that identified a number of potential sRNAs, but none have yet been characterized (44). However, Vc2 differs from all these previous examples as it is located at the 3’ end of the *P_1_*-Vc2 sRNA and *P_2_*-Vc2 sRNAs and regulates their levels in the cell by impacting RNA stability.

Stem-loop structures are common features at the 3’-end of bacterial mRNAs and have long been understood to provide protection against the 3’-5’ exonuclease activity of the RNA degradosome, which has a preference for segments of single-stranded RNA (45). The initial biochemical work on the Vc2 riboswitch provides clues about how the aptamer structure could serve a role in 3’-end stabilization. Specifically, the P1 stem of the Vc2 riboswitch is predicted to be the actor in Vc2 riboswitch function because it is the most labile region of the aptamer structure which is stabilized in the c-di-GMP-bound state, and proper closure of the P1 stem requires ligand binding (13, 19). In fact, the other two helical stems of the Vc2 aptamer undergo relatively little structural change during ligand binding because their preorganization is a requirement for formation of the c-di-GMP binding pocket (46). It is no coincidence then that the 3’-end of the *P_1_*-Vc2 sRNA is located immediately adjacent to the base of the P1 stem and that this was also the site of poly-adenylation for the sRNAs we recovered (Fig. 1). The doublet sizes of *P_1_*-Vc2 sRNA transcripts and the pattern of their degradation we observed in our rifampicin stability assay are consistent with distinct populations of 3’-tailed and non-tailed transcripts, which is a hallmark of the 3’-end degradative processes described for many regulatory RNAs in bacteria (Fig. 5A; (47)). We hypothesize that closure of the P1 stem of the Vc2 aptamer during a c-di-GMP binding event increases the stability of the *P_1_*-Vc2 sRNA by making the nucleotides at the 3’-end of the riboswitch transcript inaccessible to 3’-5’ exonucleases.

Previous work with the Vc2 riboswitch has made note of the exceptionally strong, picomolar affinity of the c-di-GMP aptamer for its ligand as measured *in vitro*, despite the observation that c-di-GMP concentrations *in vivo* reside in the nanomolar to low micromolar range (19, 48, 49). Specifically, ligand binding at the Vc2 aptamer experiences an unusually slow off-rate, so slow, in fact, that each c-di-GMP binding event should effectively be irreversible over the lifetime of the RNA transcript in the cell (19, 48). For canonical riboswitches that employ transcriptional mechanisms of regulation, the on-rate of ligand binding is the more important feature of aptamer kinetics because a switching decision must be made in the brief window of time that transcription of the expression platform is occurring (50). After initial ligand binding occurs, long-term retention of the ligand by the riboswitch aptamer is unimportant in transcriptional regulation because the RNA polymerase has long passed and a final decision about the “on” or “off” state of downstream gene expression has already been made. Biochemical study of the G20 and C92 sites of the Vc2 aptamer showed that the effect of those mutations was minimal on the c-di-GMP binding on-rate but had a great effect on the off-rate (19, 48). The slow off-rate of the Vc2 riboswitch is consistent with c-di-GMP-binding serving a continued function throughout the entire lifetime of the *P_1_*-Vc2 sRNA in the cell by maintaining the Vc2 aptamer in a structure that would make the nucleotides of the aptamer inaccessible to RNAses.

An important outstanding question of this research regards the function of the *P_1_*-Vc2 sRNA after expression. This intracellular concentration of *P_1_*-Vc2 sRNA increases directly with c-di-GMP, which suggests it may have a regulatory function at high c-di-GMP. Another interesting question is whether the *P_1_*-Vc2 sRNA and *P_2_*-Vc2 sRNA have different targets in the cell or control different phenotypes. One piece of evidence suggesting this might be the case is that the *P_2-tfoY_* promoter is repressed at the level of transcriptional induction by c-di-GMP whereas the *P_1-tfoY_* is not, leading to an increased abundance of the *P_1_*-Vc2 sRNA at high levels of c-di-GMP and decreased levels of the *P_2_*-Vc2 sRNA (10). Based on its expression pattern, we predict the *P_1_*-Vc2 sRNA could regulate traits associated with the biofilm lifestyle of *V. cholerae*.

## EXPERIMENTAL PROCEDURES

### Strains and Growth Conditions

The wild-type *V. cholerae* strain used in this study is CW2034, an El Tor biotype C6706str2 derivative containing a deletion of the *vpsL* gene (51). Use of a *vpsL-* strain facilitates accurate spectrophotometric readings during transcriptional and translational reporter assays due to decreased biofilm formation. Propagation of DNA for genetic manipulation and mating of reporter plasmids was conducted in *E. coli* S-17-*λpir* (52). All growth of bacteria was performed at 35°C in Miller LB Broth (Acumedia), or on LB agar plates, with ampicillin (100 μg/mL), kanamycin (100 μg/mL), polymyxin B (10 IU/mL), streptomycin (500 μg/mL), and isopropyl-β-D-thiogalactoside (IPTG) at 100 μM when appropriate. Liquid cultures in tubes or flasks were continuously shaken at 220 RPM; microplate cultures were continuously shaken at 150 RPM.

### Genetic Manipulations

*V. cholerae* mutant strains were constructed using allelic exchange with vectors derived from pKAS32 (53). Vectors containing mutant riboswitch alleles were generated with the QuickChange Site-Directed Mutagenesis Kit (Agilent) and mated into *V. cholerae* so as to create markerless strains with single point mutations on the genome.

### 3’-RACE

The 3’-RACE method was adapted from (54). A DNA oligonucleotide adapter was synthesized with a monophosphate at the 5’-end and an inverted thymidine base at the 3’-end (IDT). 500 pmol of adapter was ligated to 10 µg *V. cholerae* RNA at 37°C with 20 units of T4 RNA ligase (NEB) in a 20 µL reaction containing 1X T4 RNA ligase buffer, 1mM ATP, 10% DMSO, and 20 units RNasin (Promega). Ligation products were reverse transcribed into cDNA and amplified with the Access RT-PCR Kit (Promega) using a primer complementary to the 3’ adapter and a primer specific for the 5’-end sequence of transcripts initiated at the *P_1-tfoY_* promoter. The products generated were electrophoresed on an agarose gel, and DNA was excised and purified from all size ranges and recovered by TOPO TA cloning into pCR2.1 (Invitrogen).

### Northern Blotting

RNA was extracted from mid-log phase cultures using TRIzol Reagent (Invitrogen). Samples were normalized to the same concentration using spectrophotometric measurement and evaluated for ribosomal RNA integrity using a Bioanalyzer (Agilent). RNA was separated by PAGE, transferred by semi-dry blotting onto a positively-charged nylon membrane, and baked in a vacuum oven. The membrane was hybridized with probe at 65°C for at least 4 hours in ULTRAhyb buffer (Ambion) and washed three times with 0.1x SSC at 65°C. The probe was detected by chemiluminescence using the Phototope-Star Detection Kit (NEB) and autoradiographic film. Film images were digitized by scanning at high resolution with a Typhoon FLA 9500 Imager (GE Healthcare), and quantitative measurements were using the image analysis software Fiji (55).

In all, four RNA probes named probe *P_1_*, probe *P_2_*, probe *P_1+2_*, and probe Vc2 were used, which are complementary to the genomic regions −337 to −272, −272 to −191, −337 to −191, and −211 to −94, respectively, relative to the coding sequence of *tfoY* (Fig. 2A). Probes were labeled with Bio-11-UTP during *in vitro* transcription with the MAXIscript T7 Kit (Ambion) from PCR-derived DNA templates containing a consensus T7 promoter fused to the appropriate sequence.

### *in vitro* Transcription

DNA templates for in vitro transcription were generated by PCR from the vectors used for construction of *V. cholerae* genomic riboswitch mutants. Templates encompassed the genomic region from −535 nt to +120 nt relative to the *tfoY* coding sequence. The template for the *C. difficile* class-II riboswitch encompassed the region from −761 nt to −505 nt relative to the coding sequence of the gene *CD3267* of *C. difficile* 630, the same region investigated by Lee et al. 2010. Transcription reactions included 150 ng of DNA template, 2 mM DTT, 0.25mM NTPs, 0.1mM c-di-GMP when required, and 0.3 units of σ^70^-saturated *E. coli* RNA polymerase holoenzyme (Epicentre) in the transcription buffer recommended by the manufacturer. Bio-11-UTP (Ambion) was used in reactions at a concentration of 20% of total UTP to label transcripts after it had been determined not to interfere with the generation of transcription products (data not shown).

### RNA Stability Assay

200 ml cultures were grown shaking in baffled flasks after being started with a 1:10,000 dilution of overnight culture into fresh media. When the optical density reached ∼0.400-0.500, 10 mL of culture was withdrawn for a time zero reading and rifampicin was added to the culture for a final concentration of 250 µg/mL. Ten more mL of culture was removed at each subsequent time point, stabilized with 1 mL of RNA stop solution (10% phenol in 95% ethanol), and placed on ice, as described by Bernstein et al., 2002. RNA was promptly extracted from the cells using Trizol Reagent (Invitrogen) according to the manufacturer’s instructions. RNA samples were normalized, electrophoresed, blotted, and detected as described above. For quantitative analysis, a series of film exposures over multiple lengths of time were collected, and once digitized, images were only compared between images for which the film exposure was not saturated.

## Acknowledgments

This material is based in part upon work supported by the National Science Foundation under Cooperative Agreements MCB-1253684, DBI-0939454, and NIH grants GM109259 and GM110444.

## APPENDIX

**Table 1:** Strain List.

| Name | Genotype | Reference |
| --- | --- | --- |
| CW2034 | *ΔvpsL* | (51) |
| BP20 | *ΔvpsL ΔVc2* | this study |
| BP33 | *ΔvpsL Vc2* (G83U) | this study |
| BP34 | *ΔvpsL Vc2* (C92U) | this study |
| BP35 | *ΔvpsL Vc2* (G20U,C92U) | (10) |

**Table 2:**
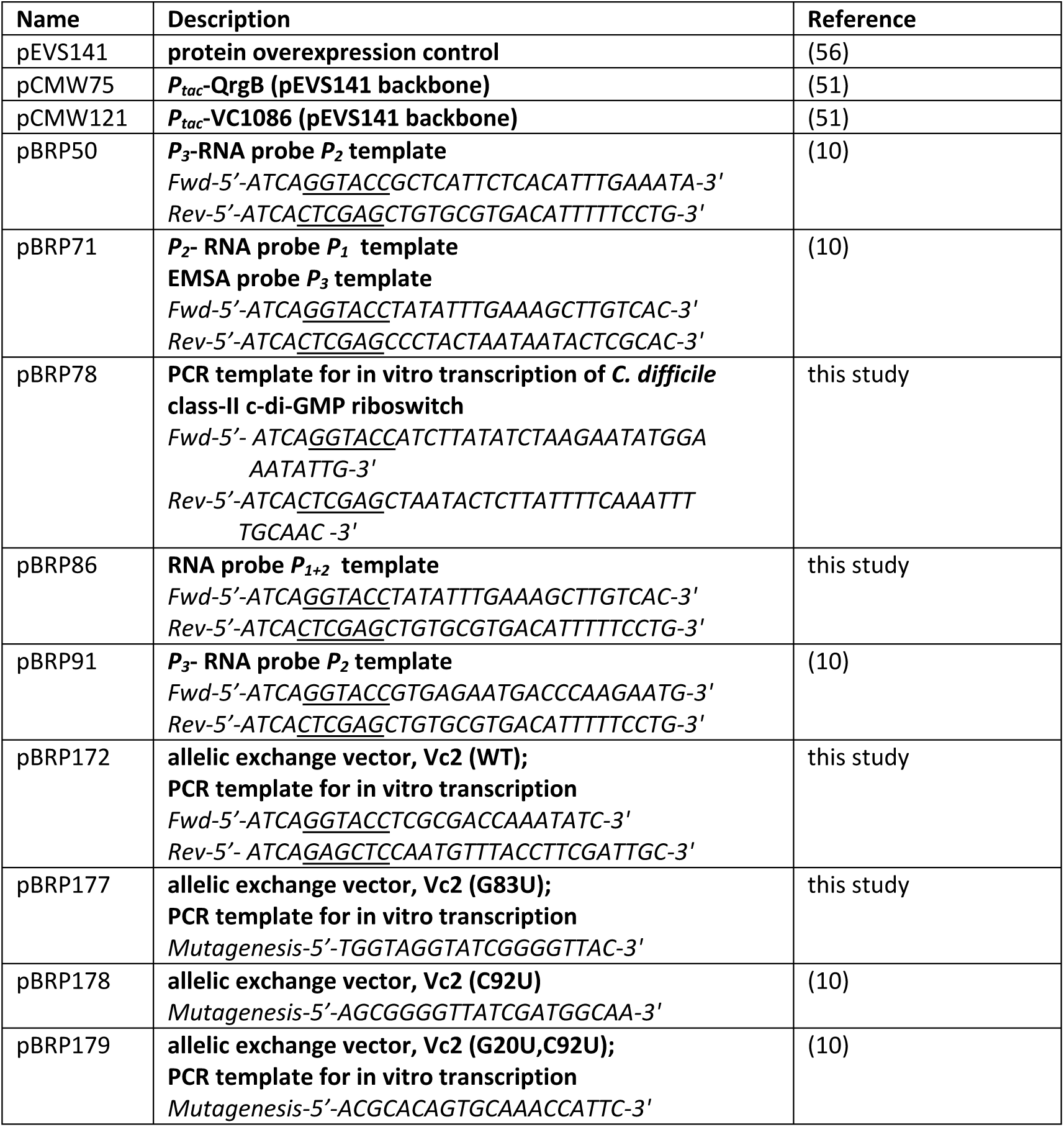
Vector and Primer List. For each vector constructed for this study, sequences are given for primers used in PCR to generate vector inserts. For primer sequences, restriction endonuclease sites are indicated by underline.

